## supplementary figures and tables for "Obesogenic diet in mice leads to inflammation and oxidative stress in the mother in association with sex-specific changes in fetal development, inflammatory markers and placental transcriptome"

### Supplementary Figure 1

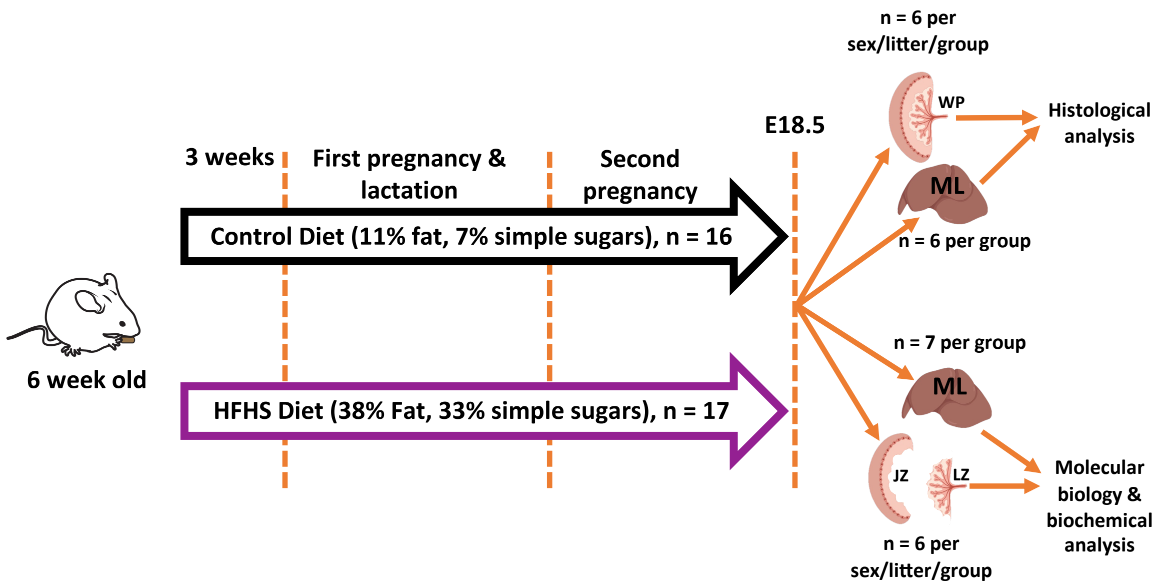


**ML: Maternal liver**
**JZ: Placental junctional zone**
**LZ: Placental labyrinth zone**
**WP: Whole placenta**

Supplementary Figure 2

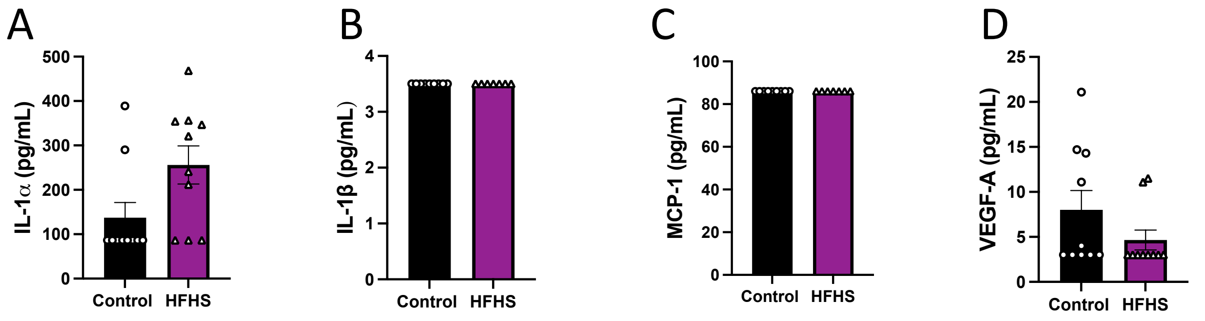

Supplementary Figure 3

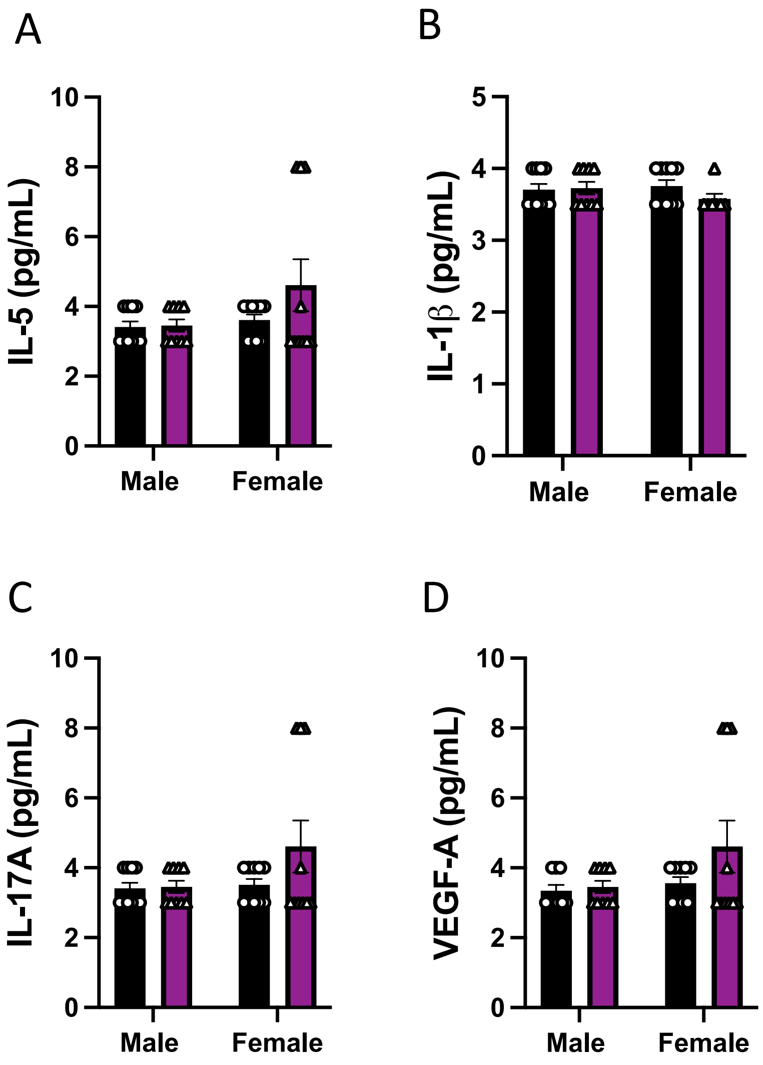

### Supplementary Figure 4

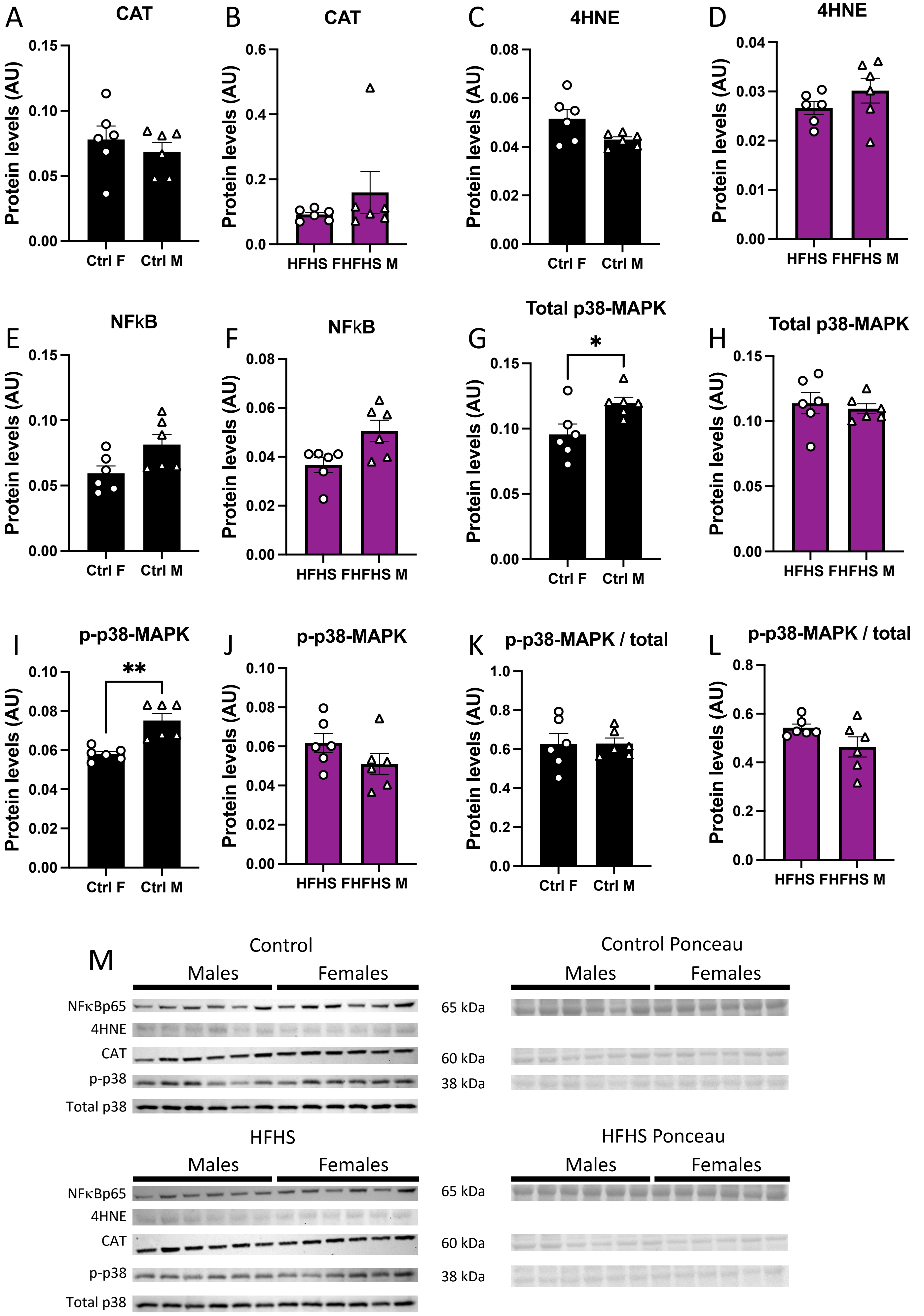

Supplementary Table 1. **DEGs related to biological process of interest – Upregulated**

| | Biological process related to: | | | | | | Altered when comparing Ctrl vs HFHS: | | |
|---|---|---|---|---|---|---|---|---|---|
| Upregulated | Immune response | Lipid or fatty acid metabolic process | Response to hypoxia & ROS | Ion transport | Nutrients transport | Angiogenesis | Overall | Females | Males |
| Cd28 | x | | | | | | Yes | No | No |
| Rasgrp1 | x | | | | | | Yes | No | Yes |
| Tgtp1 | x | | | | | | Yes | No | No |
| Chil3 | x | | | | | | Yes | Yes | Yes |
| Hamp | x | | | | | | Yes | No | Yes |
| Il1a | x | | | | | | Yes | No | Yes |
| Pla2g4 | x | | | | | | Yes | Yes | Yes |
| Oas1b | x | | | | | | No | No | Yes |
| Clec2d | x | | | | | | No | No | Yes |
| Gdap10 | x | | | | | | No | No | Yes |
| Il18bp | x | | | | | | No | No | Yes |
| Lgals4 | x | | | | | | No | No | Yes |
| Pisd-ps1 | x | | | | | | No | No | Yes |
| Snhg1 | x | | | | | | Yes | No | Yes |
| Snhg20 | x | | | | | | No | No | Yes |
| Bdh2 | | x | | | | | Yes | Yes | No |
| Apold1 | | x | x | | x | x | Yes | No | Yes |
| Cyb5r2 | | x | | | | | Yes | No | Yes |
| Gdpd3 | | x | | | | | Yes | No | No |
| Hpgds | | x | | | | | Yes | No | Yes |
| Pla2g4a | | x | | | | | Yes | Yes | Yes |
| Alms1 | | x | | | | | No | No | Yes |
| Acsbg1 | | x | | | | | Yes | No | Yes |
| Acsm3 | | x | | | | | Yes | No | No |
| Echdc2 | | x | | | | | No | No | Yes |
| Phka2 | | x | | | | | No | No | Yes |
| Nox4 | | | x | | | | Yes | No | No |
| Gucy1a2 | | | x | | | | Yes | No | No |
| Ptprn | | | x | | | | Yes | No | No |
| Hamp | | | | x | | | Yes | No | Yes |
| Rgs4 | | | | x | | | Yes | No | Yes |
| Slc10a6 | | | | x | | | Yes | No | No |
| Slc26a7 | | | | x | | | Yes | Yes | No |
| Slc6a14 | | | | x | | | Yes | Yes | No |
| Slc9b2 | | | | x | | | Yes | Yes | Yes |
| Slc30a4 | | | | x | | | No | Yes | No |
| Slc9b1 | | | | x | | | No | Yes | No |
| Gabre | | | | x | | | No | No | Yes |
| Gabrr2 | | | | x | | | No | No | Yes |
| Atg9b | | | | | x | | No | No | Yes |
| Spx | | | | | x | | No | No | Yes |
| Fgf9 | | | | | | x | Yes | No | No |

### Supplementary Table 2. **DEGs related to biological process of interest** - Downregulated

| | Biological process related to: | | | | | | Altered when comparing Ctrl vs HFHS: | | |
|---|---|---|---|---|---|---|---|---|---|
| Downregulated DEGs | Immune response | Lipid or fatty acid metabolic process | Response to hypoxia & ROS | Ion transport | Nutrients transport | Angiogenesis | Overall | Females | Males |
| Ihh | x | | | | | x | Yes | No | Yes |
| Pbk | x | | | | | | Yes | No | No |
| Ager | x | | x | | | | Yes | Yes | Yes |
| Camk4 | x | | | | | | Yes | Yes | Yes |
| Ccdc88b | x | | | | | | Yes | No | No |
| Eda | x | | | | | | Yes | Yes | Yes |
| Igha | x | | | | | | Yes | Yes | Yes |
| Jchain | x | | | | | | Yes | Yes | Yes |
| Igkc | x | | | | | | Yes | Yes | Yes |
| Jaml | x | | | | | | Yes | No | No |
| Lpo | x | | x | | | | Yes | No | No |
| Pdgfb | x | | | | | x | Yes | Yes | Yes |
| Susd4 | x | | | | | | Yes | Yes | Yes |
| Unc5cl | x | | | | | | Yes | Yes | Yes |
| Vegfd | x | | x | | | x | Yes | No | Yes |
| Cebpb | x | | | | | | No | Yes | No |
| Fkbp1b | x | | x | | | | No | Yes | No |
| 1700037H04Rik | x | | | | | | No | Yes | No |
| Thrsp | x | x | | | | | Yes | Yes | No |
| Wnt4 | x | | | | | | Yes | Yes | Yes |
| Apoe | x | x | | | x | | No | No | Yes |
| Btn1a1 | x | | | | | | No | No | Yes |
| Cx3cr1 | x | | | | | x | No | No | Yes |
| Hspa5 | x | | | | | | No | No | Yes |
| Ifi30 | x | | | | | | No | No | Yes |
| Lgals2 | x | | | | | | No | No | Yes |
| Ptgfr | x | | | | | | No | No | Yes |
| Spn | x | | | | | | No | No | Yes |
| Elovl4 | | x | | | | | Yes | No | Yes |
| Hf4a | | x | | | | | Yes | No | Yes |
| Ltc4s | | x | | | | | Yes | No | No |
| Scd2 | | x | | | | | Yes | Yes | Yes |
| Lipg | | x | | | | | No | No | Yes |
| Ccna2 | | | x | | | | Yes | No | No |
| Slc2a8 | | | x | | x | | No | Yes | No |
| Hsp90b1 | | | x | | | | No | No | Yes |
| Atp6v0e2 | | | | x | | | Yes | No | Yes |
| Kcnip3 | | | | x | | | Yes | No | No |
| Kcnip2 | | | | x | | | Yes | Yes | Yes |
| Pdzd3 | | | | x | | | Yes | No | Yes |
| Wnk2 | | | | x | | | Yes | Yes | No |
| Hpn | | | | x | | | Yes | No | Yes |
| Kcnn1 | | | | x | | | Yes | No | Yes |
| Cacna1g | | | | x | | | Yes | Yes | Yes |
| Snta1 | | | | x | | | Yes | Yes | No |
| Kcnk6 | | | | x | | | No | No | Yes |
| Trpv6 | | | | x | | | No | No | Yes |
| Anpep | | | | | | x | Yes | Yes | Yes |
| Aplnr | | | | | | x | Yes | Yes | Yes |
| Cadm4 | | | | | | x | Yes | Yes | Yes |
| Enpep | | | | | | x | Yes | No | Yes |
| Mmrn2 | | | | | | x | Yes | No | No |
| Pdgfb | | | | | | x | Yes | Yes | Yes |
| Smo | | | | | | x | Yes | No | Yes |
| Creb3l1 | | | | | | x | No | No | Yes |
| Gja5 | | | | | | x | No | No | Yes |
| Wnt7b | | | | | | x | No | No | Yes |

### Supplementary Table 3

| | | Lien et al. 2020 | | Candia et al. 2023 | |
|---|---|---|---|---|---|
| | | **LogFC** | Direction of change | LogFC | Direcction of change |
| Overal | Apoa2 | 0.4676 | Upregulated | -1.664 | Downregulated |
| | Derl3 | -0.2909 | Downregulated | -0.8214 | Downregulated |
| | Hnf4a | 0.2383 | Upregulated | -0.6725 | Downregulated |
| | Igfbp1 | 0.1945 | Upregulated | -0.8751 | Downregulated |
| | Endou | -0.2226 | Downregulated | -0.6593 | Downregulated |
| | Igfbp6 | -0.4652 | Downregulated | -0.9445 | Downregulated |
| | Scd2 | 0.3195 | Upregulated | -0.8336 | Downregulated |
| | Ccdc184 | 1.249 | Upregulated | -0.5979 | Downregulated |
| | Elovl4 | -0.2856 | Downregulated | -0.8061 | Downregulated |
| | Abtb2 | 0.6324 | Upregulated | -0.5853 | Downregulated |
| | Spata2l | -0.7153 | Downregulated | -0.6037 | Downregulated |
| | Cdhr2 | 0.4082 | Upregulated | -1.981 | Downregulated |
| | Thrsp | -0.5856 | Downregulated | -0.749 | Downregulated |
| | Cysrt1 | -0.308 | Downregulated | -0.677 | Downregulated |
| | Asb10 | 0.3591 | Upregulated | -0.6987 | Downregulated |
| | Pcp4l1 | -0.3033 | Downregulated | -0.7501 | Downregulated |
| | Unc5cl | 0.395 | Upregulated | -0.9061 | Downregulated |
| | Rarres1 | -0.2644 | Downregulated | -0.6824 | Downregulated |
| | Spink2 | 0.0994 | Upregulated | -0.7265 | Downregulated |
| | Chst8 | 0.3231 | Upregulated | -0.5981 | Downregulated |
| | Itih5l-ps | -0.3358 | Downregulated | -0.7817 | Downregulated |
| | Gm6967 | -0.4597 | Downregulated | -0.8069 | Downregulated |
| | Adora2b | 1.2272 | Upregulated | 0.5969 | Upregulated |
| | 4930412O13Rik | 0.3467 | Upregulated | 0.8344 | Upregulated |
| | Il1a | 1.6012 | Upregulated | 0.9643 | Upregulated |
| | Slc10a6 | 0.6251 | Upregulated | 0.7399 | Upregulated |
| | Ing4 | -0.2686 | Downregulated | 0.7233 | Upregulated |
| | Gdpd3 | 1.0079 | Upregulated | 2.972 | Upregulated |
| | Slc6a14 | 0.3296 | Upregulated | 0.5905 | Upregulated |
| | Rgs4 | 0.6455 | Upregulated | 0.6297 | Upregulated |
| | Chil3 | -0.5241 | Downregulated | 1.416 | Upregulated |
| | Samd9l | 1.2516 | Upregulated | 0.6415 | Upregulated |
| | Hamp | 1.1941 | Upregulated | 1.18 | Upregulated |
| | Tgtp1 | 1.321 | Upregulated | 0.9841 | Upregulated |
| | Gm15972 | -0.4264 | Downregulated | 0.6378 | Upregulated |
| | Apold1 | 0.9598 | Upregulated | 0.8594 | Upregulated |
| | Gm20627 | -0.373 | Downregulated | 0.7379 | Upregulated |
| | D5Ertd605e | -0.3257 | Downregulated | 0.6978 | Upregulated |
| Females | Scd2 | 0.3195 | Upregulated | -0.635 | Downregulated |
| | Thrsp | -0.5856 | Downregulated | -0.9754 | Downregulated |
| | Pnoc | -0.2869 | Downregulated | -2.585 | Downregulated |
| | Slc6a14 | 0.3296 | Upregulated | 0.7507 | Upregulated |
| | Chil3 | -0.5241 | Downregulated | 1.268 | Upregulated |
| Males | Ltbp1 | 0.3008 | Upregulated | -0.6296 | Downregulated |
| | Htra1 | 0.3239 | Upregulated | -0.6936 | Downregulated |
| | Slc5a1 | 0.3765 | Upregulated | -1.662 | Downregulated |
| | Slc9a5 | 0.3937 | Upregulated | 0.6846 | Upregulated |
| | Wnt7b | -0.2844 | Downregulated | -0.6233 | Downregulated |
| | Endou | -0.2226 | Downregulated | -0.6909 | Downregulated |
| | Igfbp6 | -0.4652 | Downregulated | -0.9746 | Downregulated |
| | Pgc | -0.092 | Downregulated | -0.6118 | Downregulated |
| | Zfp397 | -0.3159 | Downregulated | 0.6907 | Upregulated |
| | Cpn1 | 0.3036 | Upregulated | -1.923 | Downregulated |
| | Scd2 | 0.3195 | Upregulated | -0.9842 | Downregulated |
| | 4930412O13Rik | 0.3467 | Upregulated | 1.449 | Upregulated |
| | Il1a | 1.6012 | Upregulated | 1.345 | Upregulated |
| | Echdc2 | 0.4017 | Upregulated | 0.6956 | Upregulated |
| | Oas1b | 0.7805 | Upregulated | 0.7144 | Upregulated |
| | 1600015I10Rik | -0.5099 | Downregulated | 0.8993 | Upregulated |
| | Trpv6 | -0.2448 | Downregulated | -0.7122 | Downregulated |
| | Clec2d | 0.3596 | Upregulated | 0.6639 | Upregulated |
| | Ing4 | -0.2686 | Downregulated | 1.082 | Upregulated |
| | Syt8 | 0.2537 | Upregulated | 0.7586 | Upregulated |
| | Elovl4 | -0.2856 | Downregulated | -0.895 | Downregulated |
| | Uba7 | 0.9647 | Upregulated | 0.951 | Upregulated |
| | Ppip5k1 | 0.2791 | Upregulated | 0.7134 | Upregulated |
| | Fam208b | 0.2572 | Upregulated | 0.5851 | Upregulated |
| | Ptgdr2 | -0.5465 | Downregulated | -0.603 | Downregulated |
| | Cdhr2 | 0.4082 | Upregulated | -2.572 | Downregulated |
| | Creb3l3 | 0.3708 | Upregulated | -1.15 | Downregulated |
| | Cysrt1 | -0.308 | Downregulated | -0.6707 | Downregulated |
| | Asb10 | 0.3591 | Upregulated | -0.8802 | Downregulated |
| | Rgs4 | 0.6455 | Upregulated | 0.7965 | Upregulated |
| | A230050P20Rik | 0.6014 | Upregulated | 0.6085 | Upregulated |
| | Bace2 | 0.2868 | Upregulated | -0.7172 | Downregulated |
| | Chil3 | -0.5241 | Downregulated | 1.599 | Upregulated |
| | 2010003K11Rik | 0.1909 | Upregulated | -2.457 | Downregulated |
| | Lgals2 | 0.2674 | Upregulated | -1.132 | Downregulated |
| | Pnoc | -0.2869 | Downregulated | -1.972 | Downregulated |
| | Samd9l | 1.2516 | Upregulated | 0.8596 | Upregulated |
| | Rarres1 | -0.2644 | Downregulated | -0.7834 | Downregulated |
| | Hamp | 1.1941 | Upregulated | 1.234 | Upregulated |
| | Cx3cr1 | -0.2885 | Downregulated | -0.7852 | Downregulated |
| | Spink2 | 0.0994 | Upregulated | -0.8786 | Downregulated |
| | 2010109I03Rik | 0.5219 | Upregulated | 0.7922 | Upregulated |
| | Hsp25-ps1 | 0.8689 | Upregulated | -0.6478 | Downregulated |
| | Pisd-ps1 | -0.5627 | Downregulated | 0.6441 | Upregulated |
| | Apold1 | 0.9598 | Upregulated | 0.947 | Upregulated |
| | Gm20627 | -0.373 | Downregulated | 0.9368 | Upregulated |
| | Gdap10 | 0.318 | Upregulated | 1.103 | Upregulated |
| | Gm4673 | -0.3378 | Downregulated | 0.6196 | Upregulated |

### Supplementary Table 4. DEGs compared to an intrauterine inflammation model

| | | Lien 2020 | | Candia 2023 | |
|---|---|---|---|---|---|
| Comparison | Gene | **LogFC** | Direction of change | LogFC | Direcction of change |
| Overall | **Il1a** | 1.6012 | Upregulated | 0.9643 | Upregulated |
| | **Slc10a6** | 0.6251 | Upregulated | 0.7399 | Upregulated |
| | **Gdpd3** | 1.0079 | Upregulated | 2.972 | Upregulated |
| | **Slc6a14** | 0.3296 | Upregulated | 0.5905 | Upregulated |
| | **Rgs4** | 0.6455 | Upregulated | 0.6297 | Upregulated |
| | Chil3 | -0.5241 | Downregulated | 1.416 | Upregulated |
| | **Hamp** | 1.1941 | Upregulated | 1.18 | Upregulated |
| | **Tgtp1** | 1.321 | Upregulated | 0.9841 | Upregulated |
| | **Apold1** | 0.9598 | Upregulated | 0.8594 | Upregulated |
| | **Elovl4** | -0.2856 | Downregulated | -0.8061 | Downregulated |
| | **Thrsp** | -0.5856 | Downregulated | -0.749 | Downregulated |
| | Unc5cl | 0.395 | Upregulated | -0.9061 | Downregulated |
| | Scd2 | 0.3195 | Upregulated | -0.8336 | Downregulated |
| Females | **Slc6a14** | 0.3296 | Upregulated | 0.7507 | Upregulated |
| | Chil3 | -0.5241 | Downregulated | 1.268 | Upregulated |
| | Scd2 | 0.3195 | Upregulated | -0.635 | Downregulated |
| | **Thrsp** | -0.5856 | Downregulated | -0.9754 | Downregulated |
| Males | **Il1a** | 1.6012 | Upregulated | 1.345 | Upregulated |
| | **Echdc2** | 0.4017 | Upregulated | 0.6956 | Upregulated |
| | **Oas1b** | 0.7805 | Upregulated | 0.7144 | Upregulated |
| | **Clec2d** | 0.3596 | Upregulated | 0.6639 | Upregulated |
| | **Rgs4** | 0.6455 | Upregulated | 0.7965 | Upregulated |
| | Chil3 | -0.5241 | Downregulated | 1.599 | Upregulated |
| | **Hamp** | 1.1941 | Upregulated | 1.234 | Upregulated |
| | Pisd-ps1 | -0.5627 | Downregulated | 0.6441 | Upregulated |
| | **Apold1** | 0.9598 | Upregulated | 0.947 | Upregulated |
| | **Gdap10** | 0.318 | Upregulated | 1.103 | Upregulated |
| | **Wnt7b** | -0.2844 | Downregulated | -0.6233 | Downregulated |
| | Scd2 | 0.3195 | Upregulated | -0.9842 | Downregulated |
| | **Trpv6** | -0.2448 | Downregulated | -0.7122 | Downregulated |
| | **Elovl4** | -0.2856 | Downregulated | -0.895 | Downregulated |
| | Lgals2 | 0.2674 | Upregulated | -1.132 | Downregulated |
| | **Cx3cr1** | -0.2885 | Downregulated | -0.7852 | Downregulated |
